# Using Core Genome Alignments to Assign Bacterial Species

**DOI:** 10.1101/328021

**Authors:** Matthew Chung, James B. Munro, Julie C. Dunning Hotopp

**Author notes:** Address correspondence to Julie C. Dunning Hotopp,.

## Abstract

With the exponential increase in the number of bacterial taxa with genome sequence data, a new standardized method is needed to assign bacterial species designations using genomic data that is consistent with the classically-obtained taxonomy. This is particularly acute for unculturable obligate intracellular bacteria like those in the Rickettsiales, where classical methods like DNA-DNA hybridization cannot be used to define species. Within the Rickettsiales, species designations have been applied inconsistently, often obfuscating the relationship between organisms and the context for experimental results. In this study, we generated core genome alignments for a wide range of genera with classically defined species, including *Arcobacter*, *Caulobacter*, *Erwinia*, *Neisseria*, *Polaribacter*, *Ralstonia*, *Thermus*, as well as genera within the Rickettsiales including *Rickettsia*, *Orientia*, *Ehrlichia*, *Neoehrlichia*, *Anaplasma*, *eorickettsia*, and *Wolbachia*. A core genome alignment sequence identity (CGASI) threshold of 96.8% was found to maximize the prediction of classically-defined species. Using the CGASI cutoff, the *Wolbachia* genus can be delineated into species that differ from the currently used supergroup designations, while the *Rickettsia* genus is delineated into nine species, as opposed to the current 27 species. Additionally, we find that core genome alignments cannot be constructed between genomes belonging to different genera, establishing a bacterial genus cutoff that suggests the need to create new genera from the *Anaplasma* and *Neorickettsia*. By using core genome alignments to assign taxonomic designations, we aim to provide a high-resolution, robust method for bacterial nomenclature that is aligned with classically-obtained results.

## IMPORTANCE

With the increasing availability of genome sequences, we sought to develop and apply a robust, high-resolution method for the assignment of genera and species designations that can recapitulate classically-defined taxonomic designations. We developed genera and species cutoffs using both the length and sequence identity of core genome alignments as taxonomic criteria, respectively. These criteria were then tested on diverse bacterial genera with an emphasis on the taxonomy of organisms within the order Rickettsiales, where species designations have been applied inconsistently. Our results indicate that the *Rickettsia* have an overabundance of species designations and that there are clear demarcations of *Wolbachia* species that do not align precisely with the existing supergroup designations. Lastly, we find that the current *Anaplasma* and *Neorickettsia* genus designations are both too broad and need to be divided.

## INTRODUCTION

While acknowledging the disdain some scientists have for taxonomy, Stephen Jay Gould frequently highlighted in his writings how the classifications arising from a good taxonomy both reflects and directs our thinking, stating, “the way we order reflects the way we think. Historical changes in classification are the fossilized indicators of conceptual revolutions” (1). Historically, bacterial species delimitation relied on the phenotypic, morphological, and chemotaxonomic characterization (2–4). The 1960s saw the introduction of molecular techniques in bacterial species delimitation through the use of GC-content (5), DNA-DNA hybridization (6), and 16S rRNA sequencing (7, 8). Currently, databases like SILVA (9) and Greengenes (10) use 16S rRNA sequencing to identify bacteria. However, 16S rRNA sequencing often fails to separate closely-related taxa, and its utility for species-level identification is questionable (10–12). Multilocus sequence analysis (MLSA) has also been used to determine species (13), as has phylogenetic analysis of both rRNA and protein-coding genes (3, 14, 15). Non-genomic mass-spectrometry-based approaches, in which expressed proteins and peptides are characterized, provide complementary data to phenotypic and genomic species delimitations (16, 17) and are used in clinical microbiology laboratories. However, DNA-DNA hybridization (DDH) remains the “gold standard” of defining bacterial species (18, 19), despite its inability to address non-culturable organisms and the intensive labor involved that limits its applicability. A new genome-based bacterial species definition is attractive given the increasing availability of bacterial genomes, rapid sequencing improvements with decreasing sequencing costs, and data standards and databases that enable data sharing.

Average nucleotide identity (ANI) and *in silico* DDH were developed as genomic era tools that allow for bacterial classification with a high correlation to results obtained using wet lab DDH, while bypassing the associated difficulties (20–22). For ANI calculations, the genome of the query organism is split into 1 kbp fragments, which are then searched against the whole genome of a reference organism. The average sequence identity of all matches having >60% overall sequence identity over >70% of their length is defined as the ANI between the two organisms (18). *In silico* DDH, also referred to as digital DDH (dDDH), uses the sequence similarity of conserved regions between the genomes of interests, such as high scoring segment pairs (HSPs) or maximally unique matches (MUMs) (23), to calculate genome-to-genome distances. These distances are converted to a dDDH value, which is intended to be analogous to DDH values obtained using traditional laboratory methods. There are three formulas for calculating dDDH values between two genomes using either (1) the length of all HSPs divided by the total genome length, (2) the sum of all identities found in HSPs divided by the overall HSP length, and (3) the sum of all identities found in HSPs divided by the total genome length, with the second formula being recommended for assigning species designations for draft genomes (24, 25). However, these methods do not report the total length of fragments that match the reference genome, and problems arise when only a small number of fragments are unknowingly used.

Recent phylogenomic analyses have shifted towards using the core proteome, a concatenated alignment constructed using the amino acid sequences of genes shared between the organisms of interest (26). However, differences in annotation that affect gene calls can add an unnecessary variable when deriving evolutionary relationships. We propose the use of a nucleotide core genome alignment, constructed using all collinear genomic regions free of rearrangements (27), to infer phylogenomic relationships. Using the length of the core genome alignment along with a sequence similarity matrix we aim to assign taxonomic designations at the genus, species, and strain levels. A core genome alignment-based method provides advantages to its protein-based counterpart in that it is of a higher resolution, independent of annotation, transparent with respect to the data used in the calculations, and very amenable to data sharing and deposition in data repositories.

The Rickettsiales are an order within the Alphaproteobacteria is composed of exclusively obligate, intracellular bacteria where classic DNA-DNA hybridization is not possible and bacterial taxonomy is uneven, with each of the genera having its own criteria for assigning genus and species designations. Within the Rickettsiales, there are three major families: the Anaplasmataceae, Midichloriaceae, and Rickettsiaceae, with is an abundance of genomic data being available for genera within the Anaplasmataceae and Rickettsiaceae. However, species definitions in these families are inconsistent, and organisms in the *Wolbachia* genus lack community-supported species designations, instead relying on a system of supergroup designations.

The last reorganization of the Rickettsiaceae taxonomy occurred in 2001 (28), a time when there were <300 sequenced bacterial genomes (29). As of 2014, there have been more than 14,000 genomes sequenced and with this increase in available genomic information, more informed decisions can be made with regards to taxonomic classification (29). By using core genome alignments, we can condense whole genomes into positions shared between the input genomes and use sequence identity to infer phylogenomic relationships. In this study, we aim to both establish guidelines for delineating bacterial genus and species boundaries that are consistent with classically-defined genus and species designations and apply these guidelines to organisms in the Rickettsiales, including *Rickettsia*, *Orientia*, *Ehrlichia*, *Neoehrlichia*, *Anaplasma*, *Neorickettsia*, and *Wolbachia*.

## RESULTS

### Assessment of genus designations

Core genome alignments were generated using the whole genome aligner Mugsy with both complete and draft genome sequences assembled in ≤100 contigs (27). The Mugsy algorithm uses NUCmer to identify locally colinear blocks (LCBs) between the input genomes, where LCBs are shared colinear genomic regions free of rearrangements. LCBs found in all organisms are selected and filtered to include only positions present in all genomes. The resulting core genome alignment is used to generate a sequence similarity matrix.

Core genome alignments were constructed for the *Arcobacter*, *Caulobacter*, *Erwinia*, *Neisseria*, *Polaribacter*, *Ralstonia*, and *Thermus* genera. Each of the resultant core genome alignments were ≥0.33 Mbp in size and represented ≥13.6% of the average input genome sizes (Table 1, **Supplementary Table 1)**, suggesting that the technique is applicable to a wide range of classically defined bacterial taxa. Additionally, core genome alignments were successfully constructed from the genomes of four of the six Rickettsiales genera, including 69 *Rickettsia* genomes, 3 *Orientia* genomes, 16 *Ehrlichia* genomes, and 23 *Wolbachia* genomes (Table 1). Each of these four core genome alignments are >0.87 Mbp in length and contain >10% of the average size of the input genomes.

**Table 1:**
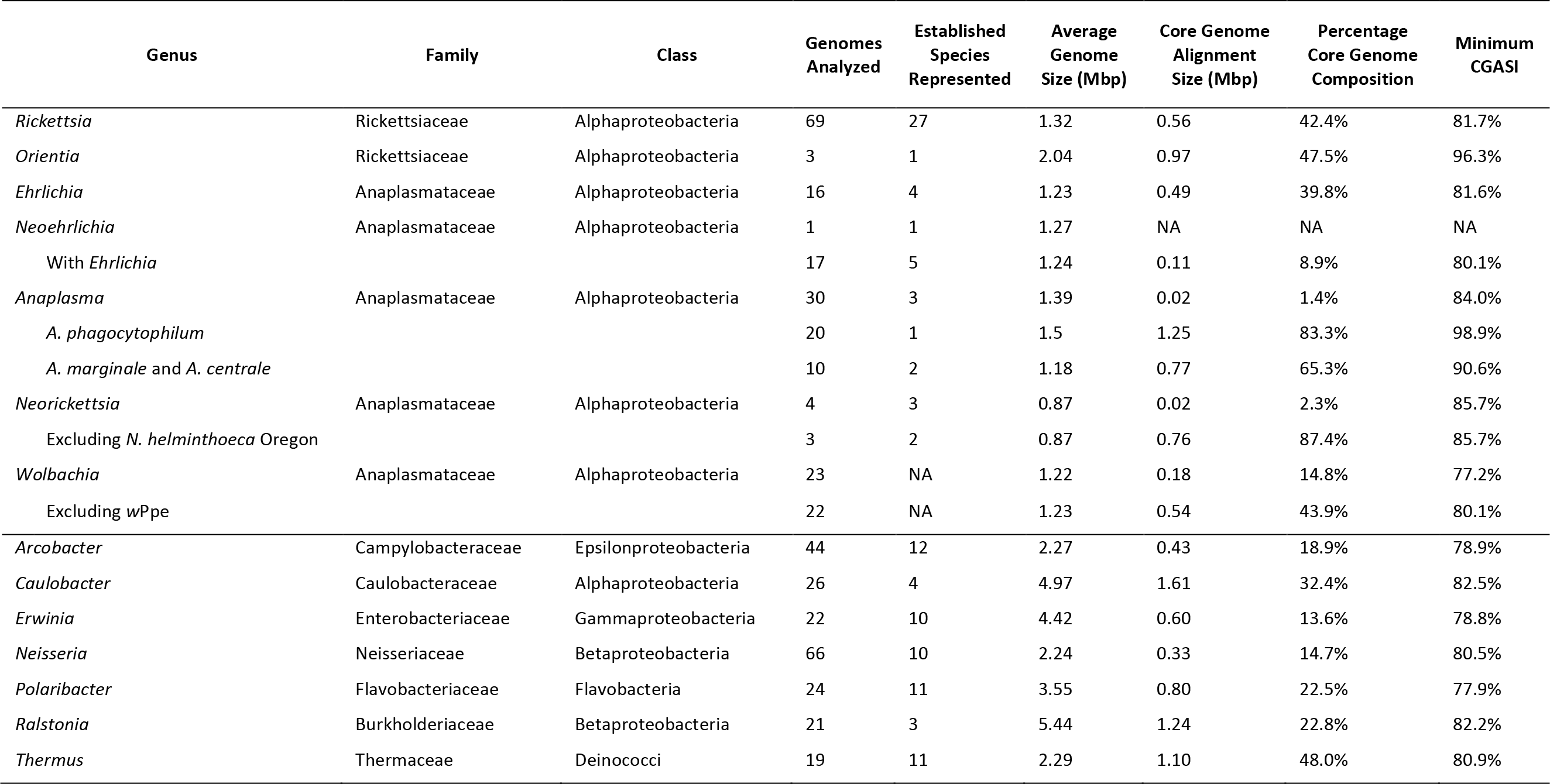
Rickettsiales genome and core genome alignment statistics.

We found our initial core genome alignments for *Anaplasma* and *Neorickettsia* to be considerably shorter, both being ~20 kbp and accounting for ≤2.3% of the average input genome sizes. We believe this reflects that the genomes within these two genera are too broad and need further refinement. Using input genomes from different genera to construct a core genome alignment yields an alignment of an insufficient size to accurately represent the evolutionary distances between the input genomes. For example, when the genome of *Neoehrlichia lotoris* is supplemented to the genomes used to create the 0.49 Mbp *Ehrlichia* core genome alignment, the resultant core genome alignment is 0.11 Mbp, representing only 8.9% of the average input genome size compared to the prior 39.8%. Therefore, we used subsets of species to test whether the *Anaplasma* and *Neorickettsia* genera are too broadly defined. A core genome alignment generated using only the 20 *A. phagocytophilum* genomes in the 30 *Anaplasma* genome set is 1.25 Mbp and represents 83.3% of the average input genome size, while a core genome alignment of the remaining 10 *Anaplasma* genomes is 0.77 Mbp, 65.3% of the average genome input size. This result suggests that the *Anaplasma* genus should be split into two separate genera. Similarly, when the genome of *N. helminthoeca* Oregon is excluded from the *Neorickettsia* core genome alignment, a 0.77 Mbp *Neorickettsia* core genome alignment is generated which represents 87.4% of the average input genome size, suggesting *N. helminthoeca* Oregon is not of the same genus as the other three *Neorickettsia* genome. For the remainder of the manuscript, these genus reclassifications are used. Given these collective results we recommend that the genus classification level can be defined as a group of genomes that together will yield a core genome alignment that represents ≥10% of the average input genome sizes.

### Advantages of nucleotide alignments over protein alignments for bacterial species analyses

While core protein alignments are increasingly used for phylogenetics, a core nucleotide alignment should have more phylogenetically informative positions in the absence of substitution saturation, yielding a greater potential for phylogenetic signal. Nucleotide-based analyses outperform amino acid-based analyses in terms of resolution, branch support, and congruence with independent evidence (30, 31) and outperform amino-acid based analyses at all time scales (32). A core protein alignment generated from 152 genes shared between the ten complete *Wolbachia* genomes contains 16,241 parsimony-informative positions while the core nucleotide alignment contains 124,074 such positions, indicating a 10-fold increase in potentially informative positions (Figure 1). Substitution saturation can negatively impact nucleotide-based phylogenetic distance measurements relative to protein-based phylogenetic methods. However, for each core genome alignment, when the uncorrected pairwise genetic distances are plotted against the model-corrected distances, linear relationships are observed for all alignments (*r*^*2*^>0.995), indicating substitution saturation is not hindering the ability of the core genome alignments to represent evolutionary relationships (33) **(Supplementary Figure 1)**. Given our results for numerous taxa, we expect this result to be broadly applicable across bacterial species. Furthermore, maximum likelihood (ML) trees from the core protein alignment and the core nucleotide alignment are quite similar for branch length as well as topology, except for the wNo branch (Figure 1), which is indicative of a longstanding problem with resolution in the *Wolbachia* phylogeny (34–37).

**Figure 1:**
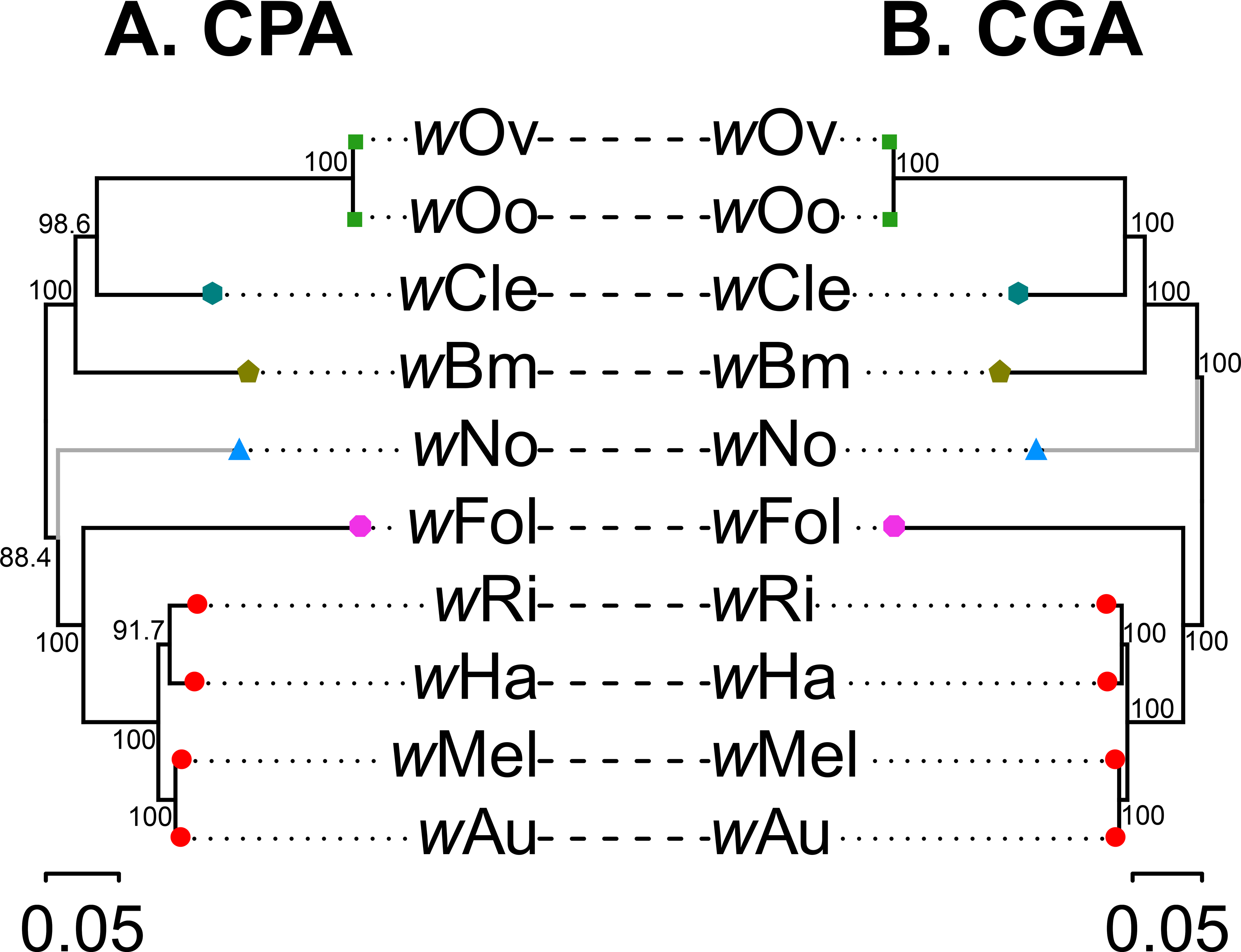
Comparison of phylogenomic trees generated using the core nucleotide and core protein alignments for ten complete *Wolbachia* genomes. Two phylogenomic trees were generated from the genomes of ten *Wolbachia* strains using **(A)** a core protein alignment (CPA) containing 152 genes present in only one copy in all ten genomes and **(B)** a core genome alignment (CNA) with members of the *Wolbachia* supergroups A (•), B (▴), C (◾), D 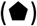, and F 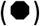 being represented. Shapes of the same color indicate that the multiple genomes are of the same species as determined using our determined CGASI cutoff of ≥96.8%. The single difference in topology is denoted in grey, otherwise the trees are largely similar in both topology and branch lengths, despite the core protein alignment being 77,868 bp long with 16,241 parsimony-informative positions while the core nucleotide alignment is 579,495 bp long with 124,074 such positions.

### Identifying a CGASI cutoff for species delineation

Within the *Rickettsia*, *Orientia*, *Ehrlichia*, *Anaplasma*, *Neorickettsia*, *Wolbachia*, *Arcobacter*, *Caulobacter*, *Erwinia*, *Neisseria*, *Polaribacter*, *Ralstonia*, and *Thermus* genome subsets, ANI, dDDH, and CGASI values were calculated for 7,264 pairwise genome comparisons **(Supplementary Table 2)**, of which 601 are between members of the same species. The ANI and CGASI follow a second-degree polynomial relationship (*r*^*2*^ = 0.977) (Figure 2A). Using this model, the ANI species cutoff of ≥95% is analogous to a CGASI cutoff of ≥96.8%. The dDDH and CGASI follow a third-degree polynomial model (*r*^*2*^ = 0.978), with a dDDH of 70% being equivalent to a CGASI of 97.6%, indicating the dDDH species cutoff is generally more stringent than the ANI species cutoff (Figure 2B).

**Figure 2:**
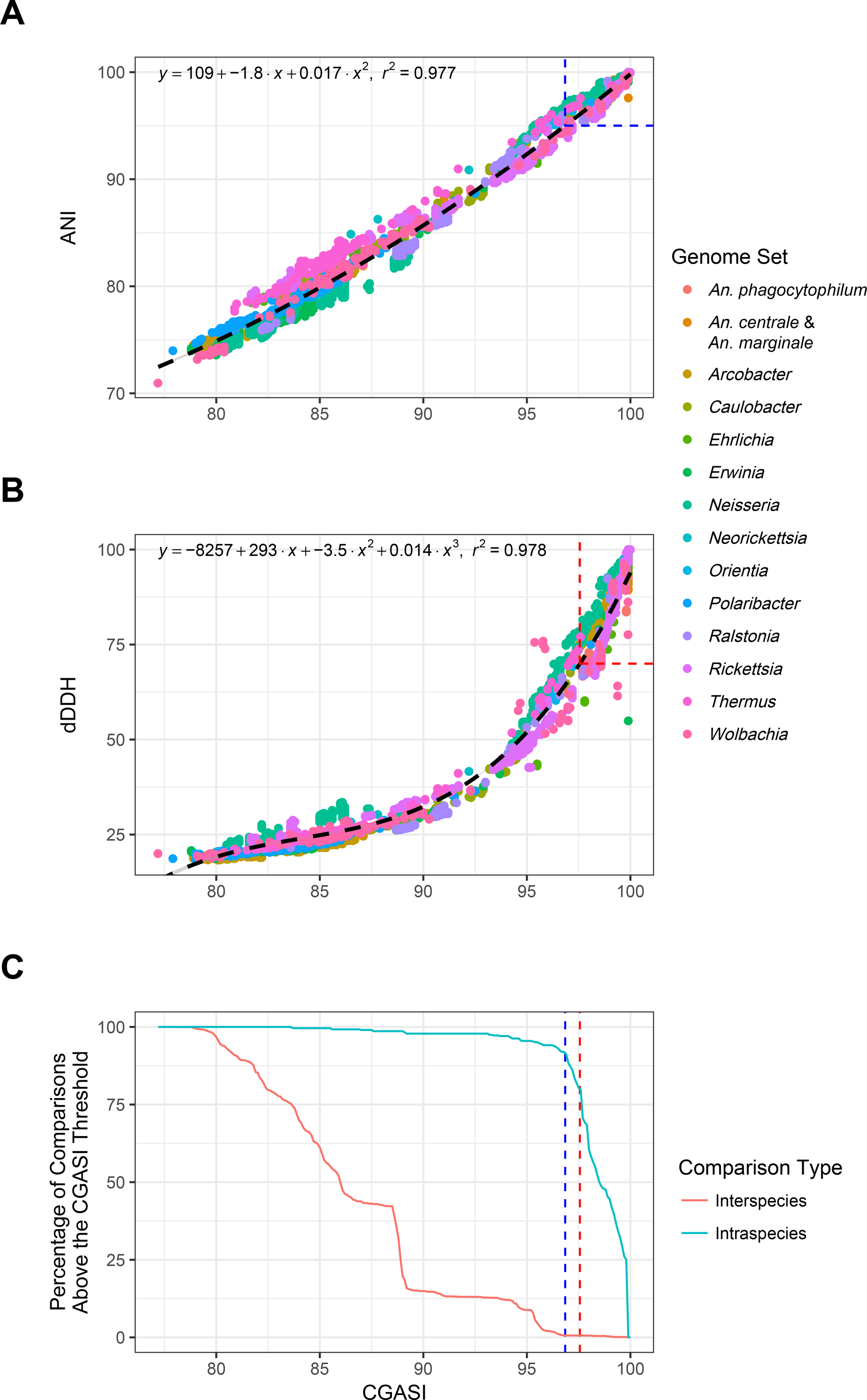
ANI, dDDH, and CGASI correlation analysis. CGASI, ANI and dDDH values were calculated for 7,264 pairwise comparisons of genomes within the same genus for *Rickettsia*, *Orientia*, *Ehrlichia*, *Anaplasma*, *Neorickettsia*, *Wolbachia*, *Caulobacter*, *Erwinia*, *Neisseria*, *Polaribacter*, *Ralstonia*, and *Thermus*. **(A)** The CGASI and ANI values for the interspecies comparisons follow a second-degree polynomial model (*r*^*2*^ = 0.977) with the ANI species cutoff of ≥95% being equivalent to a CGASI of 96.8%, indicated by the blue dotted box. **(B)** The CGASI and dDDH values for all pairwise comparisons follow a third-degree polynomial model (*r*^*2*^ = 0.978) with the dDDH species cutoff of ≥70% being equivalent to a CGASI of 97.6%, indicated by the red dotted box. **(C)** To identify the optimal CGASI cutoff to use when classifying species, for each increment of the CGASI species cutoff, plotted on the x-axis, the percentage of intraspecies and interspecies comparisons correctly assigned was determined based on classically defined species designations. The ideal cutoff should maximize the prediction of classically defined species for both interspecies and intraspecies comparisons. The ANI-equivalent CGASI species cutoff is represented by the blue dotted line while the dDDH-equivalent CGASI species cutoff is represented by the red dotted line.

The ideal CGASI threshold for species delineation would maximize the prediction of classically defined species, neither creating nor destroying the majority of the classically defined species. Therefore, all pairwise comparisons were classified as either intraspecies, between genomes with the same classically defined species designation, or interspecies, between genomes within the same genus but with different classically defined species designations. Every possible CGASI threshold value was then tested for the ability to recapitulate these classically-defined taxonomic classifications (Figure 2C). In all cases, an abnormally high number of interspecies *Rickettsia* comparisons were found above both the established ANI and dDDH species thresholds consistent with previous observations that guidelines for establishing novel *Rickettsia* species are too lax (38), and as such they were excluded from this specific analysis. Below a CGASI of 97%, classically defined species begin to be separated while organisms classically defined as different species begin to be collapsed. This coincides with the above calculated ANI-equivalent threshold but differs from the above calculated dDDH-equivalent (Figure 2C). The dDDH-equivalent CGASI threshold of 97.6% failed to predict the classically defined taxa from 100 intraspecies comparisons **(Supplementary Table 3)** while the ANI-equivalent CGASI threshold of 96.8%failed to predict the classically defined taxa from 41 intraspecies comparisons **(Supplementary Table 4)**. Given these results, we selected the ANI-equivalent CGASI value of ≥96.8% to further analyze these taxa, which results in our recommendation of specific taxonomic changes within the Rickettsiaceae, including the *Rickettsia*, *Orientia*, *Anaplasma*, *Ehrlichia*, *Neorickettsia*, and *Wolbachia*.

### Rickettsiaceae phylogenomic analyses

#### Rickettsia

The Rickettsiaceae family includes two genera, the *Rickettsia* and the *Orientia*, and while both genera are obligate intracellular bacteria, *Rickettsia* genomes have undergone more reductive evolution, having a genome size ranging from 1.1-1.5 Mbp (39) compared to the 2.0-2.2 Mbp size of the *Orientia* genome (40). Of the Rickettsiales, the *Rickettsia* genus contains the greatest number of sequenced genomes and named species, containing 69 genome assemblies in ≤100 contigs representing 27 unique species. *Rickettsia* genomes are currently classified based on the Fournier criteria, an MLST approach established in 2003 based on the sequence similarity of five conserved genes: the 16S rRNA, citrate synthase (*gltA*), and three surface-exposed protein antigens (*ompA, ompB*, and gene D) (41). To be considered a *Rickettsia* species, an isolate must have a sequence similarity of ≥98.1% 16S rRNA and ≥86.5% *gltA* to at least one preexisting *Rickettsia* species. Within the *Rickettsia*, using *ompA*, *ompB*, and gene D sequence similarities, the Fournier criteria also support the further classification of *Rickettsia* species into three groups: the typhus group, the spotted fever group (SFG), and the ancestral group (41). However, the Fournier criteria has not yet been amended to classify the more recently established transitional *Rickettsia* group (42), indicating a need to update the *Rickettsia* taxonomic scheme.

A total of 69 *Rickettsia* genomes representative of 27 different established species were used for ANI, dDDH, and core genome alignment analyses, and regardless of the method used, a major reclassification is justified (Figure 3, **Supplementary Table 1)**. A core genome alignment constructed using the 69 *Rickettsia* species genomes yielded a core genome alignment size of ~0.56 Mbp, 42.4% of the lengths of the input *Rickettsia* genomes (Table 1). Within the SFG *Rickettsia*, the CGASI between any two genomes is ≥98.2%, well within the proposed CGASI species cutoff of ≥96.8% (Figure 3), while the CGASI is ≤97.2% in the ancestral and typhus groups. If a CGASI cutoff of 96.8% is used to reclassify the *Rickettsia* species, all but two of the SFG *Rickettsia* genomes would be classified as the same species (Figure 3), with the two remaining SFG *Rickettsia* genomes, *R. monacensis* IrR Munich and *Rickettsia* sp. Humboldt, being designated as the same species. This is consistent with ANI as well, while dDDH yields conflicting results (Figure 3). For the transitional group *Rickettsia, R. akari* and *R. australis* would be collapsed into a single species due to having CGASI values of 97.2% with one another. Similarly, *R. asembonensis*, *R. felis*, and *R. hoogstraalii*, all classified as transitional group *Rickettsia*, would be collapsed into another species, all having CGASI values of 97.2% with one another. This is consistent with a phylogenomic tree generated from the *Rickettsia* core genome alignments, where the SFG *Rickettsia* genomes have far less sequence divergence compared to the rest of the *Rickettsia* genomes (Figure 3).

**Figure 3:**
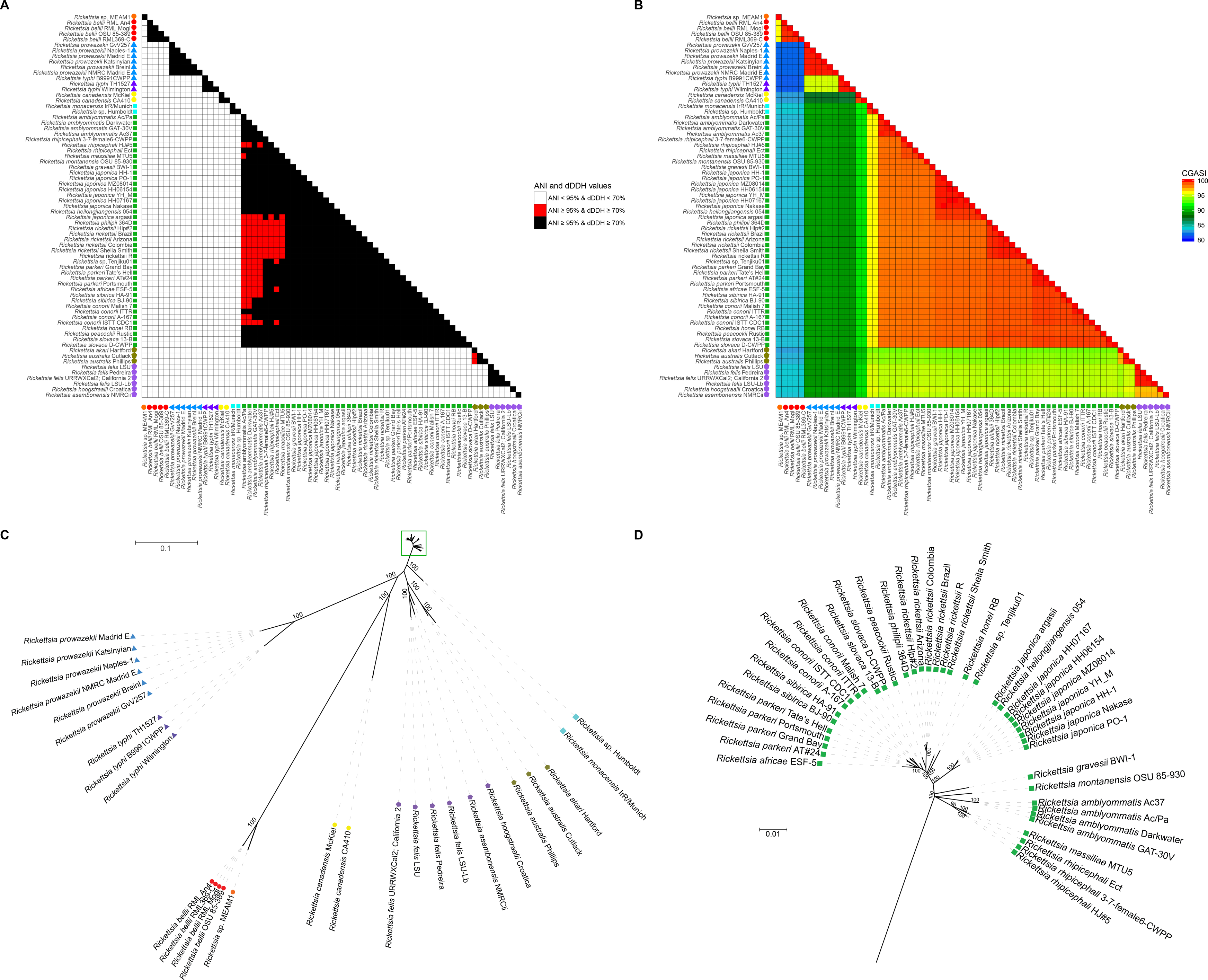
Analysis of the ANI, dDDH, and CGASI values of 69 *Rickettsia* genomes. For 69 *Rickettsia* genomes, *(A)* the ANI and dDDH values and *(B)* CGASI values were calculated for each genome comparison and color-coded to illustrate the results with respect to cutoffs of ANI ≥95% and dDDH ≥70%. The color of the shape represents species designations as determined by a CGASI cutoff ≥96.8%. **(C)** A ML phylogenomic tree with 1000 bootstraps was generated using the core genome with bootstrap support values placed next to their corresponding nodes. **(D)** The relationships in the green box on Panel C cannot be adequately visualized at the necessary scale, so they are illustrated separately with a different scale. In all panels, the shape next to each *Rickettsia* genome represents whether the genome originates from an ancestral (•), transitional 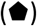, typhus group (▴), or spotted fever group (◾) *Rickettsia* species.

#### Orientia

The organisms within *Orientia* have no standardized criteria to define novel species, with *Orientia chuto* being defined as a novel species based on geographical location and the phylogenetic clustering of the 16S rRNA and two protein coding genes that encode for serine protease *htrA* (47 kDa gene) and an outer membrane protein (56-kDa gene), respectively (43). There are far fewer high-quality *Orientia* genomes, which is partly due to the large number of repeat elements found in *Orientia* genomes, with the genome of *Orientia tsutsugamushi* being the most highly repetitive sequenced bacterial genome to date with ~42% of its genome being comprised of short repetitive sequences and transposable elements (44).

The *Orientia* core genome alignment was constructed using three *O.tsutusgamushi* genomes and is 0.97 Mbp in size, ~47.6% of the average input genome size, with CGASI values ranging from 96.3-97.2% **(Supplementary Figure 2)**. Reclassifying the *Orientia* genomes using a CGASI species cutoff of 96.8% would result in *O. tsutsugamushi* Gillliam and *O.tsutsugamushi* Ikeda being classified as a separate species from *O. tsutsugamushi* Boryong **(Supplementary Figure 2)**. In this case, this reclassification would not be consistent with recommendations from using ANI or dDDH, which is likely in part due to an imperfect correlation with the ANI and dDDH species cutoffs (18). In this case, we suspect that ANI is strongly influenced by the large number of repeats in the genome due to ANI calculations being based off the sequence identity of 1 kbp query genome fragments. In comparison, we do not anticipate that whole genome alignments would be confounded by the repeats. While the LCBs may be fragmented by the repeats, creating smaller syntenic blocks, the non-phylogenetically informative repeats are eliminated from an LCB-based analysis.

### Anaplasmataceae phylogenomic analyses

#### Ehrlichia

Within the Anaplasmataceae, species designations are frequently assigned based on sequence similarity and clustering patterns from phylogenetic analyses generated using the sequences of specific genes, like the 16S rRNA, *groEL*, and *gltA* (45). As an example, the species designations for *Ehrlichia khabarensis* and *Ehrlichia ornithorhynchi* are justified based on having a lower sequence similarity for the 16S rRNA, *groEL*, and *gltA* below the maximum similarity that differentiates other *Ehrlichia* species (45–47).

A core genome alignment constructed using 16 *Ehrlichia* genomes, representative of four defined species, yields a 0.49 Mbp alignment, which equates to 39.8% of the average *Ehrlichia* genome size (1.25 Mbp) (Table 1). Using a CGASI species cutoff of 96.8%, the *Ehrlichia chaffeensis* and *Ehrlichia ruminatium* genomes were recovered as monophyletic and well supported species, which is consistent with dDDH and ANI (Figure 4). The two *Ehrlichia muris* and *Ehrlichia* sp. Wisconsin_h genomes have CGASI values >97.8%, indicating the three genomes represent one species, which is consistent with ANI, but not dDDH (Figure 4). The genomes of *Ehrlichia* sp. HF and *E. canis* Jake do not have CGASI values ≥96.8% with any other species, confirming their status as individual species, consistent with dDDH and ANI (Figure 4). This is consistent with an *Ehrlichia* phylogenomic tree (Figure 4).

**Figure 4:**
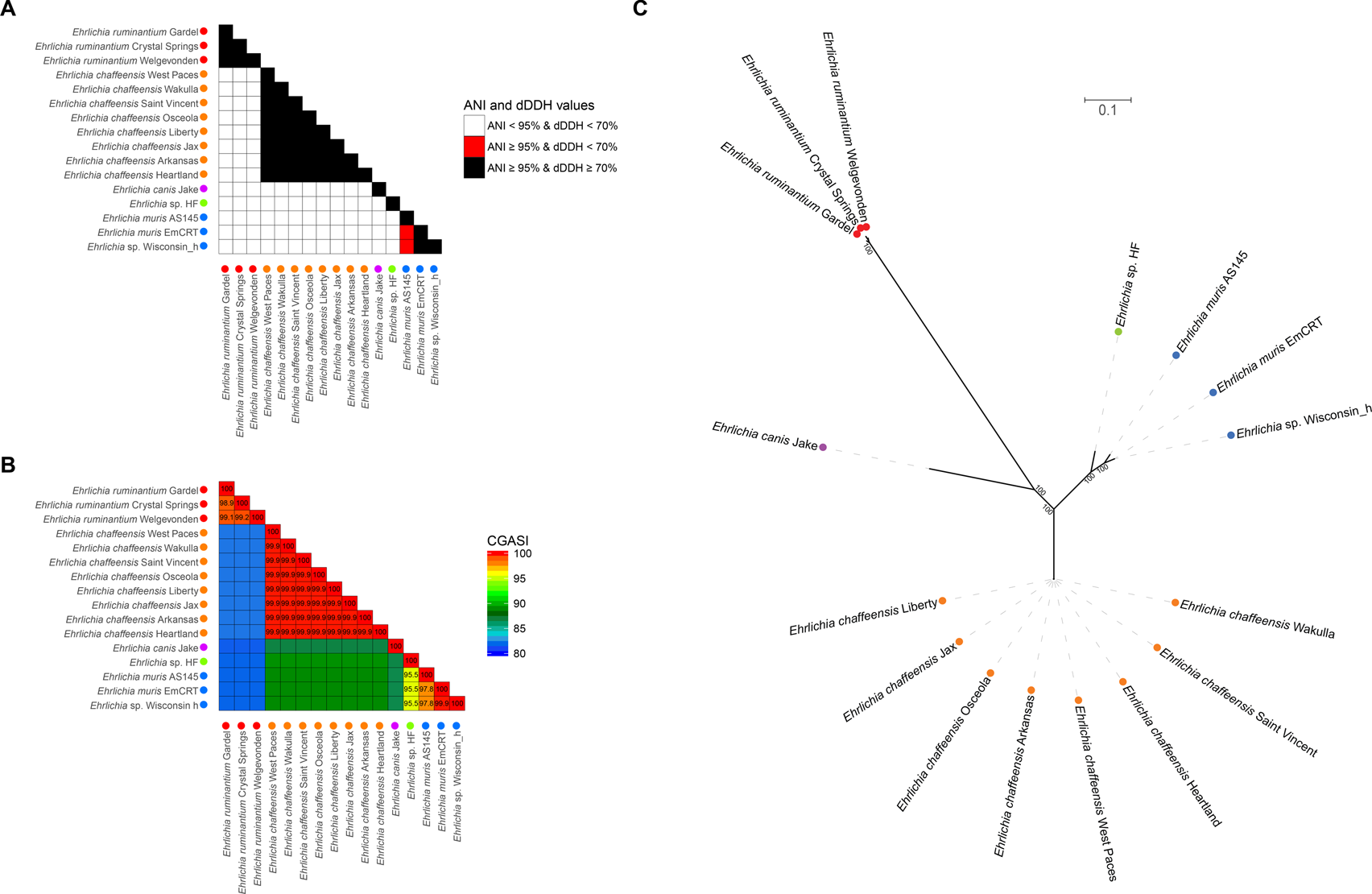
Analysis of the ANI, dDDH, and CGASI values of 16 *Ehrlichia* genomes. For 16 *Ehrlichia* genomes, **(A)** the ANI and dDDH values and **(B)** CGASI values were calculated for each genome comparison and color-coded to illustrate the results with respect to cutoffs of ANI ≥95% and dDDH ≥70%. The color of the shape represents species designations as determined by a CGASI cutoff ≥96.8%. **(C)** A ML phylogenomic tree with 1000 bootstraps was generated using the core genome with bootstrap support values placed next to their corresponding nodes.

#### Anaplasma

A novel *Anaplasma* species is currently defined based on phylogenetic analyses involving the 16S rRNA, *gltA*, and *groEL*, with a new species having a lower sequence identity and a divergent phylogenetic position relative to established *Anaplasma* species (48–51). A core genome alignment constructed using 30 *Anaplasma* genomes yielded a 20 kbp alignment, 1.4% of the average input genome size. As noted before, such low values are indicative of more than one genus represented in the taxa included in the analysis. Thus, CGASI analyses for the *Anaplasma* were done on two *Anaplasma* genome subsets, one containing the twenty *Anaplasma phagocytophilum* genomes and the other containing nine *Anaplasma marginale* and one *Anaplama centrale* genomes (Table 1).

A 1.25 Mbp core genome alignment, consisting of 39.8% of the average input genome size, was constructed using twenty *A. phagocytophilum* genomes that all have CGASI values of ≥96.8%, supporting their designation as members of a single species (Figure 5). This is supported by ANI, but dDDH again yields conflicting results (Figure 5). The core genome alignment generated using the remaining ten *Anaplasma* genomes yields a 0.77 Mbp core genome alignment, 65.3% of the average input genome size. The *A. centrale* genome has CGASI values ranging from 90.7-91.0% when compared to the nine *A. marginale* genomes, supporting *A. centrale* as a separate species from *A. marginale*, consistent with existing taxonomy and with the ANI and dDDH species cutoffs (Figure 5).

**Figure 5:**
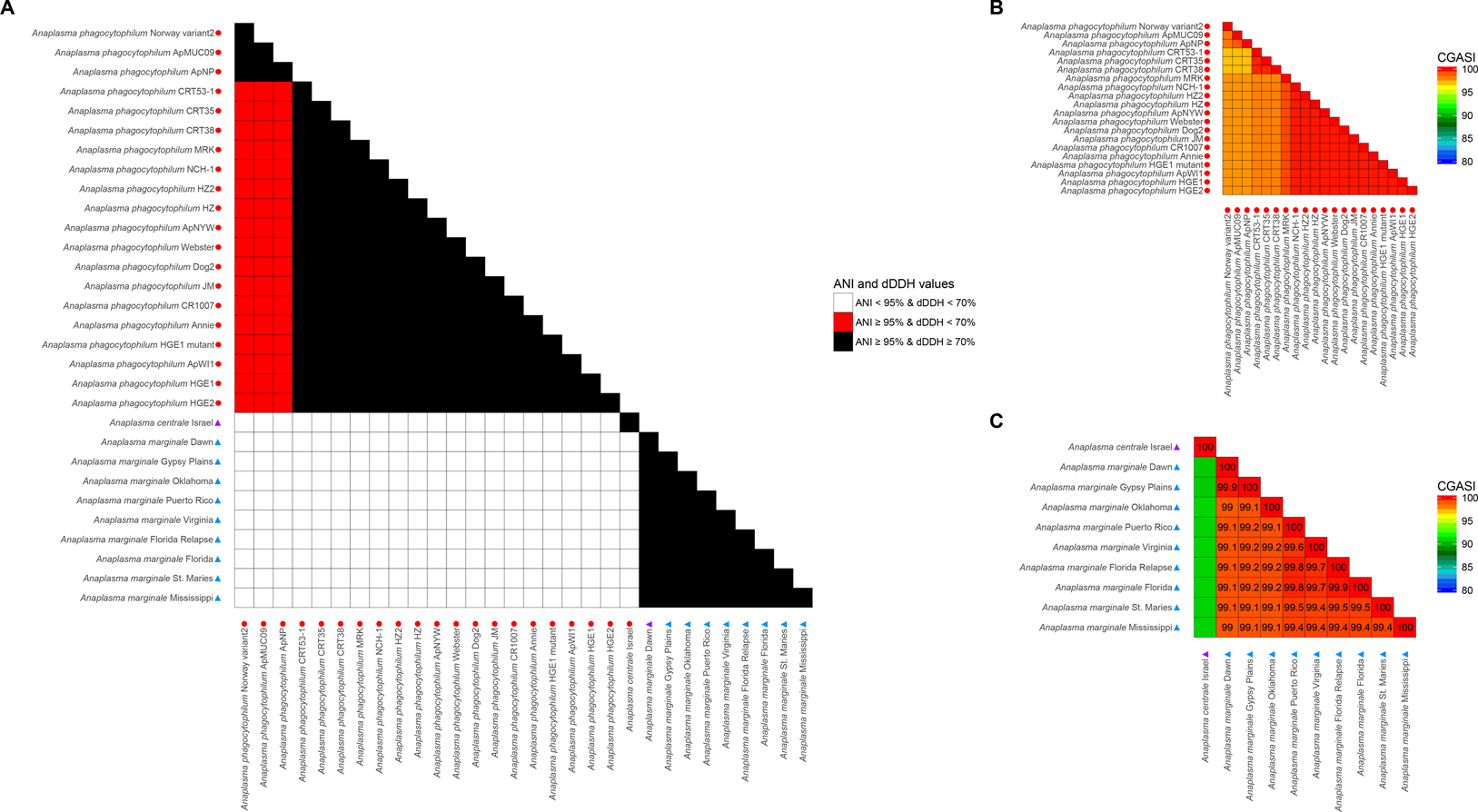
Analysis of the ANI, dDDH, and CGASI values of 30 *Anaplasma* genomes. **(A)** For 30 *Anaplasma* genomes, the ANI and dDDH values were calculated for each genome comparison and color-coded to illustrate the results with respect to cutoffs of ANI ≥95% and dDDH ≥70%. When attempting to construct a core genome alignment using all 30 *Anaplasma* genomes, only a 20-kbp alignment was generated, accounting for <1% of the average *Anaplasma* genome size. Therefore, CGASI values were calculated after the *Anaplasma* genomes were split into two subsets containing **(B)** twenty *A. phagocytophilum* genomes, and **(C)** the remaining ten *Anaplasma* genomes. Two phylogenomic trees were generated using the core genome alignments from **(D)** 20 *A. phagocytophilum* genomes and **(E)** the remaining ten *Anaplasma* genomes. In all panels, the shape next to each genome denotes genus designations as determined by CGASI while the color of the shape denotes the species as defined by a CGASI cutoff of ≥96.85%.

#### Neorickettsia

The *Neorickettsia* genus contains four genome assemblies: *Neorickettsia helminthoeca* Oregon, *Neorickettsia risticii* Illinois, *Neorickettsia sennetsu* Miyayama, and *Neorickettsia* sp. 179522. The genus was first established in 1954 with the discovery of *N. helminthoeca* (52) and in 2001, *N. risticii* and *N sennetsu*, both initially classified as *Ehrlichia* strains, were added to the *Neorickettsia* based on phylogenetic analyses of 16S rRNA and *groESL* (28). A core genome alignment constructed using all four *Neorickettsia* genomes yields a 20 kbp alignment, 2.3% of the average input genome size. When excluding the genome of *N. helminthoeca* Oregon, the three remaining *Neorickettsia* genomes form a core genome alignment of 0.76 Mbp in size, 87.4% of the average input genome size, indicating *N. risticii* Illinois, *N. sennetsu* Miyayama, and *Neorickettsia* sp. 179522 are three distinct species within the same genus while *N. helminthoeca* Oregon is of a separate genus **(Supplementary Figure 3)**. When assessing the *Neorickettsia* species designations using ANI and dDDH cutoffs, the four *Neorickettsia* genomes can only be determined to be different species, as the two techniques are unable to delineate phylogenomic relationships at the genus level.

#### Wolbachia

The current *Wolbachia* classification system currently lacks traditional species designations and instead groups organisms by supergroup designations using an MLST system consisting of 450-500 bp internal fragments of five genes, *gatB*, *coxA*, *hcpA*, *ftsZ*, and *fbpA* (53). A core genome alignment generated using 23 *Wolbachia* genomes yields a 0.18 Mbp alignment, amounting to 14.8% of the average input genome size. The CGASI values between the supergroup A *Wolbachia*, apart from *w*Inc SM, have CGASI values ≥96.8 (Figure 6). The genome of *w*Inc SM has CGASI values ranging from 94.6%-95.9% when compared to other supergroup A *Wolbachia*, indicating that *w*Inc SM is a different species. This is also supported by ANI and, with a few exceptions, dDDH. However, a phylogenomic tree generated using the *Wolbachia* core genome alignment indicates *w*Inc SM is nested within the other supergroup A *Wolbachia* taxa, a clade with 100% bootstrap support (Figure 6). The ANI, dDDH, and CGASI values of *w*Inc SM are likely underestimated due to >10% of the genome having ambiguous nucleotide positions, leading to large penalties in sequence identity scores, indicating that these methods may not be suited for genomes with large numbers of ambiguous positions. Among the filarial *Wolbachia* supergroups C and D, the CGASI cutoff of 96.8% would split each of the traditionally recognized supergroups into two groups each, also supported by ANI and dDDH. While *w*Oo and *w*Ov would be the same species, *w*Di Pavia should be considered a different species. Similarly, *w*Bm and *w*Wb would be considered the same species, while *w*Ls should be designated as a separate species. The *w*Cle, *w*Fol, and *w*Ppe endosymbionts from supergroups E, F, and L, respectively, would all be considered distinct species using CGASI, ANI, or dDDH.

**Figure 6:**
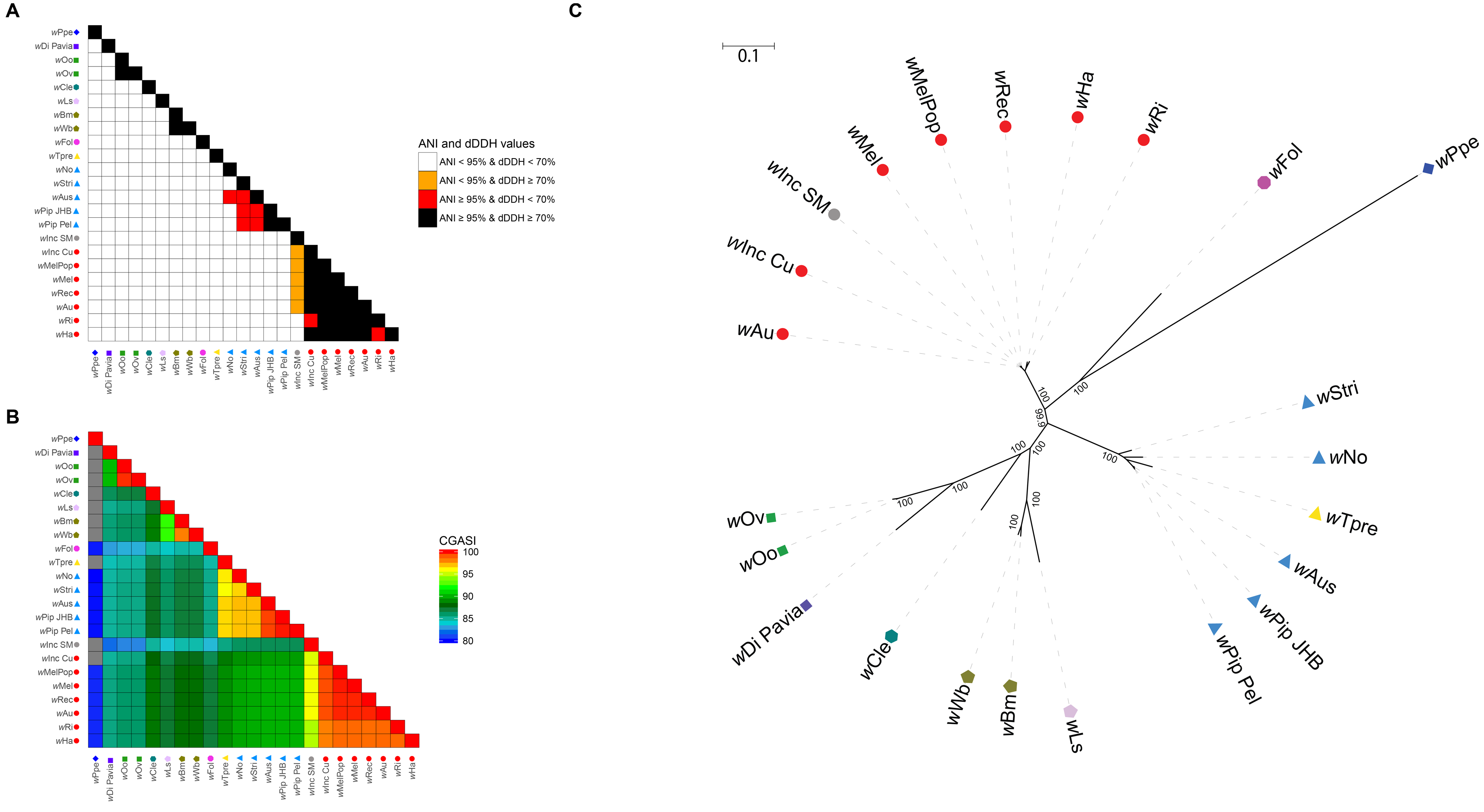
Analysis of the ANI, dDDH values of 23 *Wolbachia* genomes and CGASI values of 22 *Wolbachia* genomes. For 23 *Wolbachia* genomes, **(A)** The ANI and dDDH values and **(B)** CGASI values were calculated for each genome comparison and color-coded to illustrate the results with respect to cutoffs of ANI ≥95% and dDDH ≥70%. **(C)** A ML phylogenomic tree was generated using the *Wolbachia* core genome alignment constructed using 22 *Wolbachia* genomes. The species designations in supergroup B show the CGASI-designated species cluster of *w*Pip_Pel, *w*Pip_JHB, *w*Aus, and *w*Stri to be polyphyletic, indicating the need for amendments to the CGASI species criteria. In all panels, the shape denotes the current supergroups A (•), B (▲), C (◾), D 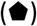, E 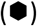, F 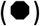 and L (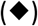) and the color denotes the species as defined by a CGASI cutoff of ≥96.85%.

The results between CGASI, ANI, and dDDH in supergroup B are discordant. Five of the six supergroup B *Wolbachia* genomes, all except for *w*Tpre, have CGASI values ≥96.8% when compared to one of the other five, indicating the five genomes are of the same species. However, despite *w*Tpre being considered a different species if the CGASI cutoff of 96.8% is used, a phylogenomic tree constructed using the core genome alignment shows *w*Tpre to be nested within the supergroup B *Wolbachia* (Figure 6). If *w*Tpre is designated as a different species, this would create a paraphyletic clade in the supergroup B *Wolbachia*, suggesting that *w*Tpre is not a different species, but rather should be included with the other supergroup B *Wolbachia* as a single species. When assessing the ANI and dDDH values between the supergroup B *Wolbachia*, only *w*Pip JHB and *w*Pip Pel share both ANI values ≥95% and dDDH values ≥70% with one another (Figure 6). The remaining three supergroup B *Wolbachia*, *w*No, *w*Stri, and *w*Aus have ANI values ≥95% with some of the other supergroup B *Wolbachia* while not having a dDDH ≥70% with any other genome, suggesting each supergroup B genome may be a separate species. Currently, the underlying basis for the differences in CGASI, ANI, and dDDH in supergroup B *Wolbachia* endosymbionts are not apparent. Given tradition, we would recommend caution and label all supergroup B *Wolbachia* as a single species with a potential re-evaluation in the future.

### Other taxa

Overall, there were 41 pairwise comparisons of organisms across diverse taxa including *Arcobacter*, *Caulobacter*, *Neisseria*, *Orientia*, *Ralstonia*, and *Thermus* that are classically defined as the same species but would be classified as distinct species when assessed using our suggested CGASI threshold **(Supplementary Table 4)**. Additionally, there were 782 pairwise comparisons of organisms classically defined as different species that this threshold suggests should be the same species, of which only 10 were not in the genus *Rickettsia*, instead belonging to the *Arcobacter* and *Caulobacter* **(Supplementary Table 5)**. This suggests that using CGASI with this threshold yields robust results. However, as seen with the supergroup B *Wolbachia* results, using strictly a nucleotide identity-based threshold can lead to paraphyletic groups, such that examination with a phylogenomic tree constructed using the core genome alignment is recommended, as was also observed for *Neisseria*.

A 0.33 Mbp core genome alignment was generated from 66 *Neisseria* genomes, accounting for 14.7% of the average input genome size. The core genome alignment largely supports the classically defined species including *N. meningitidis*, *N. gonorrhoeae*, *N. weaveri*, and *N. lactamica*, although one *N. lactamica* isolate, *N. lactamica* 338.rep1_NLAC, appears to be inaccurately assigned **(Supplementary Figure 4)**. The CGASI values suggest each *N. elongata* isolate is a distinct species, as are the *N. flavescens* isolates. Like what was observed in the *Wolbachia*, a paraphyletic clade is observed when using a CGASI cutoff of 98.6% with the genome of *Neisseria* sp. HMSC061B04 being nested within a clade of two *N. mucosa* genomes, *N. lactamica* 338.rep1_NLAC, and several unnamed *Neisseria* taxa, while not having a CGASI ≥98.6% when compared to any other *Neisseria* genome **(Supplementary Figure 4)**. This highlights that while sequence identity cutoffs derived using nucleotide-based pairwise comparisons are important when delineating species, analyses using phylogenomic trees can aid in resolving instances near the CGASI species cutoff to ensure that paraphyletic clades do not occur. The two results are quite complimentary with the core genome alignment being amenable to ML-based phylogenetic approaches.

## DISCUSSION

As noted with the development of MLST (54), sequence data has the advantages of being incredibly standardized and portable. While MLST methods allow for insight at the sub-species levels of taxonomic classifications, and are heavily relied upon during infectious disease outbreaks, they do not provide sufficient resolution at the species level. Increased resolution is required when inferring evolutionary relationships and can be obtained through the construction of whole genome alignments that maximize the number of evolutionarily informative positions.

By generating core genome alignments for different genera, we sought to identify a universal, high-resolution method for species classification. Using the core genome alignment, a set of genomes can be analyzed based on both the length and sequence identity of its core genome alignment. The length of the core genome alignment, which reflects the ability for the genomes to be aligned, is informative in delineating genera. Meanwhile, the pairwise comparisons of core genome alignment sequence identity can be used to delineate species, which can in turn be further validated with a phylogenetic analysis that can be used to resolve paraphyletic clades.

A current issue of the Mugsy aligner is the inability for the software to scale, being only able to computationally handle subsets of at most ~80 genomes. However, for heavily sequenced genera such as the *Rickettsia*, future species assignments for novel genomes do not require constructing a core genome alignment from every available genome to infer evolutionary relationships. Instead, we recommend that for each genus, the relevant experts in the community establish and curate a set of trusted genomes with at least one representative of each named species that should be used for constructing core genome alignments. After an initial assessment, refinement could then be made using a core genome alignment with many more genomes from closely related species.

Through this work we have identified criteria that largely reconstruct classical species definitions in a method that is transparent and portable. In the course of this work we have identified modifications that need to be made to the species and genus designations of a number of organisms, particularly within the Rickettsiales. While we have identified these instances, we recommend that any changes in nomenclature be addressed by collaborative teams of experts in the respective communities.

## MATERIALS AND METHODS

### Core genome alignments

Genomes used in the taxonomic analyses were downloaded from NCBI GenBank (55). OrthoANIu v1.2 (56) and USEARCH v6.1.544 (57) were used for the average nucleotide identity calculations. GGDC v2.1 (ggdc.dsmz.de) (25) paired with the recommended BLAST+ alignment tool (58) was used to calculate dDDH values. For analyses in this paper, all dDDH calculations were performed using dDDH formula 2 due to the usage of draft genomes in taxonomic analyses (24). Core genome alignments were generated for each of the genome subsets using Mugsy v1.2 (27) and MOTHUR v1.22 (59). Sequence identity matrices for the core genome alignments were created using BioEdit v7.2.5 (brownlab.mbio.ncsu.edu/JWB/papers/1999Hall1.pdf). ML phylogenomic trees with 1000 bootstraps were calculated for each core genome alignment using IQ-TREE v1.6.2 (60) paired with ModelFinder (61) to select the best model of evolution and UFBoot2 (62) for fast bootstrap approximation. Trees were visualized and annotated using iTOL v4.1.1 (63). Construction of neighbor-network trees were done using the R packages ape (64) and phangorn (65).

### Core protein alignments

Orthologs between complete genomes of the same species were determined using FastOrtho, a reimplementation of OrthoMCL (66) that identifies orthologs using all by all BLAST searches. The amino acid sequences of proteins present in all organisms in only one copy were aligned using MAAFT v7.313 (67). For every protein alignment, the best model of evolution was identified using ModelFinder (61) and phylogenomic trees were constructed using an edge-proportional partition model (68) with IQ-TREE v1.6.2 (60) and UFBoot2 (62) for fast bootstrap approximation. Comparative analyses of core protein and core nucleotide alignment trees were done using the R packages ape (64) and phangorn (65).

## ACKNOWLEDGMENTS

We would like to thank Dr. Joseph J. Gillespie for helpful discussions and encouragement. This work was funded by the National Institute of Allergy and Infectious Diseases (U19AI110820).

## SUPPLEMENTARY FIGURE LEGENDS

**Supplementary Figure 1: Assessing substitution saturation for core genome alignments**

For each of the 14 core genome alignments that comprise ≥10% of the average input genome size, the uncorrected genetic distance between each of the members was plotted against the TN69-model corrected genetic distance. The red-line represents the best-fit line for each data set while the black, dotted-line represents the identity line (y=x). In all cases, the relationship between the two distances are linear (*r*^*2*^ >0.995), indicating little substitution saturation in the core genome alignments.

**Supplementary Figure 2: Analysis of the ANI, dDDH, and CGASI values of three *Orientia* genomes**

For 3 *Orientia tsutsugamushi* genomes, **(A)** the ANI and dDDH values and **(B)** CGASI values were calculated for each genome comparison and color-coded to illustrate the results with respect to cutoffs of ANI ≥95% and dDDH ≥70%. Circles of the sample colors next to the names of each genome indicate members of the same species as defined by a CGASI ≥96.8%.

**Supplementary Figure 3: Analysis of the ANI, dDDH, and CGASI of 4 *Neorickettsia* genomes**

**(a)** For the 4 *Neorickettsia* genomes, the ANI and dDDH values were calculated for each genome comparison and color-coded to illustrate the results with respect to cutoffs of ANI ≥95% and dDDH ≥70%. **(b)** CGASI values were calculated and are illustrated using a core genome alignment that could only be constructed using 3 of the *Neorickettsia* genomes, excluding *N. helminthoeca* Oregon. Circles of the sample colors next to the names of each genome indicate members of the same species as defined by a CGASI ≥96.8%.

**Supplementary Figure 4: Analysis of the ANI, dDDH, and CGASI of 66 *Neisseria* genomes**

For the 66 *Neisseria* genomes, **(A)** the ANI and dDDH values and **(B)** CGASI values were calculated for each genome comparison and color-coded to illustrate the results with respect to cutoffs of ANI ≥95% and dDDH ≥70%. **(c)** A ML phylogenomic tree was generated using 1,000 bootstraps with the 0.33 Mbp *Neisseria* core genome alignment. Circles of the sample colors next to the names of each genome indicate members of the same species as defined by a CGASI ≥96.8

### SUPPLEMENTARY TABLES

**Supplementary Table 1: Genomes used for ANI, dDDH, and CGASI analysis**

**Supplementary Table 2: ANI, dDDH, and CGASI values for 7,264 interspecies comparisons of the *Rickettsia*, *Orientia*, *Ehrlichia*, *Anaplasma*, *Neorickettsia*, *Wolbachia*, *Arcobacter*, *Caulobacter*, *Erwinia*, *Neisseria*, *Polaribacter*, *Ralstonia*, and *Thermus* genera**

**Supplementary Table 3: ANI, dDDH, and CGASI values for 100 intraspecies comparisons with a CGASI <97.6%**

**Supplementary Table 4: ANI, dDDH, and CGASI values for 41 intraspecies comparisons with a CGASI <96.8%**

**Supplementary Table 5: ANI, dDDH, and CGASI values for 10 non-*Rickettsia* interspecies comparisons determined to be intraspecies using the ANI-derived CGASI cutoff of 96.8%**

